# Programmable transport of micro- and nanoparticles by *Paramecium caudatum*

**DOI:** 10.1101/231092

**Authors:** Richard Mayne, Jack Morgan, Neil Phillips, James Whiting, Andrew Adamatzky

## Abstract

We exploit chemo- and galvanotactic behaviour of *Paramecium caudatum* to design a hybrid device that allows for controlled uptake, transport and deposition of environmental micro- and nanoparticulates in an aqueous medium. Manipulation of these objects is specific, programmable and parallel. We demonstrate how device operation and output interpretation may be automated via a DIY low-cost fluorescence spectrometer, driven by a microprocessor board. The applications of the device presented range from collection and detoxification of environmental contaminants (e.g. nanoparticles), to micromixing, to natural expressions of computer logic.

---

The class of protistic freshwater organisms known as the ‘ciliates’ achieve locomotion, feeding and environmental sensing via the functions of cell-surface organelles known as ‘motile cilia’ (hereafter ‘cilia’). Certain varieties of metazoan epithelia, such as the columnar epithelium found in the upper respiratory tract or fallopian tubes of humans, also possess cilia. A ciliated cell may possess thousands of these hair-like organelles which achieve their purposes via a rhythmic whip-like beating motion, which creates fluid currents in surrounding fluid media. The emergent properties exhibited by collective ciliary motion, which are thought to be coordinated solely by local interactions [1, 2, 3, 4, 5, 6, 7, 8, 9, 10], have long-since been the focus of research for harnessing, mimicking and emulating for a range of biomedical and engineering uses [11, 12, 13, 14, 15, 16, 17, 18, 19, 20, 21, 22].

Artificial cilia arrays have been considered in the context of self-cleaning or anti-fouling surfaces; following these lines of thought we developed a concept of programmable intake and transport of micro-objects by cilia arrays inspired by the ciliate *Paramecium caudatum*. Such arrays were capable of orientation and transportation of various geometrical shapes [23, 24] or objects [25]. Computer models on this concept have been partially confirmed in laboratory experiments [26]. Although manipulating single micro-particles on the surface of a live ciliate is an experimentally challenging task, we anticipate value in the prospect of large scale transfer of volumes of micro-particles via controlled intake of the particle by ciliates, movement of the ciliates with particle, and programmable release of the particles.

Previously we proposed and studied in experimental laboratory conditions transfer of substances with slime mould [27], where the slime mould was stimulated to intake food colouring and propagate along the route determined by spatial configurations of sources of attractants and repellents. We a adopt similar strategy in our experiments with *P. caudatum* with the following objectives:

1. Collection of micro-scale objects from specific locations by the organisms via internalisation and intracellular carriage, ideally with some form of discrimination between objects.
2. Object transport to specific areas.
3. Flexible transport (dynamic reprogrammability of specific operations).
4. Controlled retention and deposition of ingested material.
5. Parallelism and multitasking (manipulation of multiple varieties of object and ability to perform a range of operations).
6. Modular construction and scalability. Although the capabilities of *P. caudatum* regarding their interactions with various particulates was covered thoroughly in historical literature [28, 29, 30], it was not until recently that these organisms have been characterised as doing useful ‘work’ in terms more amenable to quantitative descriptions. in our previous work, we have described *P. caudatum* interactions with environmentally-dispersed particulates as sorting (differential manipulation based on sensorial input) operations [26] and orchestrated (by humans) transport of environmental contaminants [31], as per criterion 1 in the above list.

## 1. Methods

### 1.1. P. caudatum culture

*P. caudatum* were cultivated in a modified Chalkley’s medium enriched with 10 g of dried alfalfa and 40 grains of wheat per litre at room temperature in non-sterile conditions. Cultures were exposed to a day and night cycle but were kept out of direct sunlight. Organisms were harvested in log growth phase by centrifugation at 400 × G prior to being rinsed and resuspended in dechlorinated tap water (DTW).

### 1.2. Experimental environment

The standard experimental environment (EE) (Fig. 1) used consisted of two polyethylene cuvettes measuring 12 × 12 × 44 mm, affixed to the base of a glass microscope slide with epoxy resin (Araldite, Huntsman, USA). The tubes were linked at a point 1.5 mm above their base by flexible PTFE tubing, OD 3.0 mm ID 1.0 mm length 5 mm, which were affixed to the cuvettes with epoxy. The length of the connecting tube was chosen to be short enough to reduce operation time and make the rate of diffusion proportional to the length of each experiment (see Appendix A.2). Prior to each experiment, both chambers (A, B) were filled with 2.0 ml of fresh culture medium. Care was taken to ensure that fluid levels were equal in both cuvettes and that the linking tube did not become air-locked. Whenever quantities of fluid containing cells or particles were added to a chamber, an equal quantity of fluid (DTW unless otherwise stated) was added to the other chamber simultaneously in order to prevent fluid transfer resulting from pressure changes.

**Figure 1:**
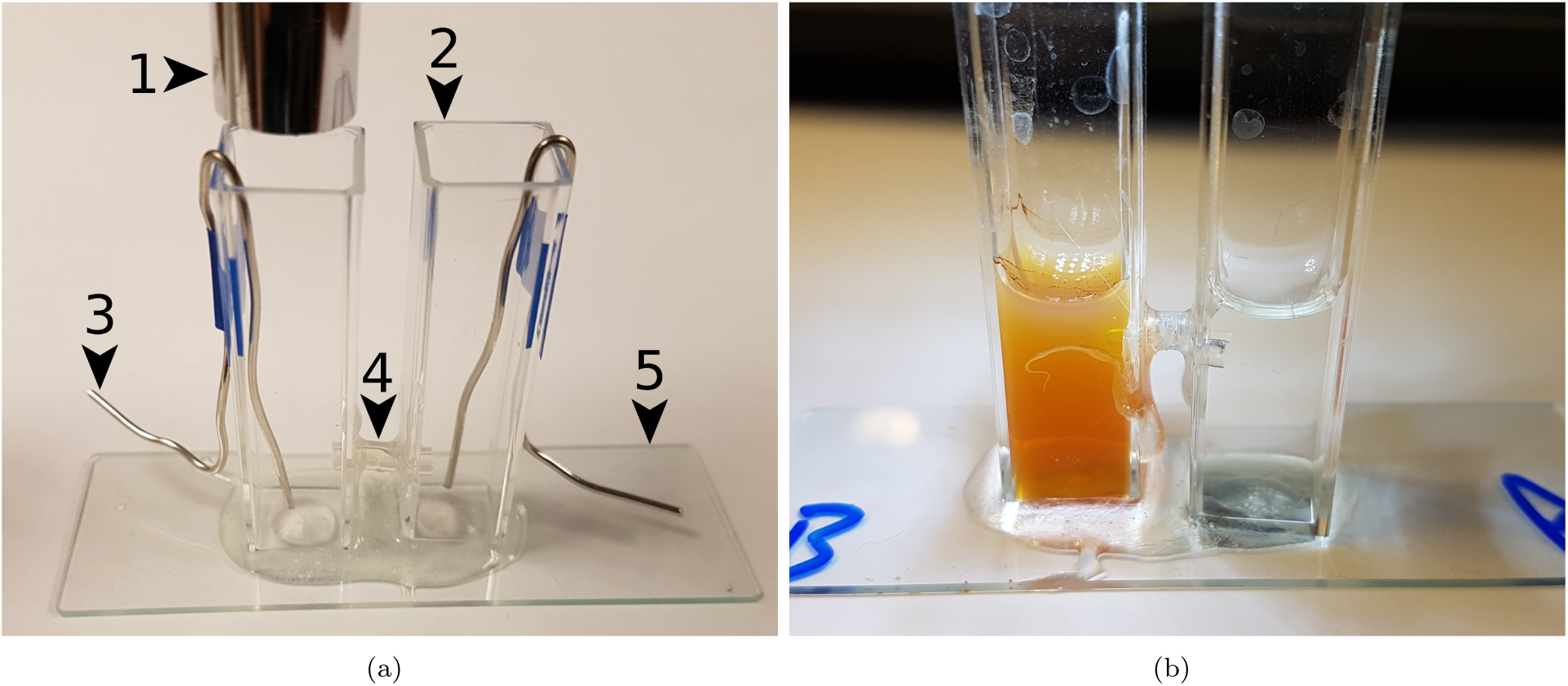
Photographs of the experimental environment. (a) Showing principle and optional components. (1) Microscope, optional. (2) Cuvette. (3) Pt electrode, optional. (4) Link tube. (5) Base, glass microscope slide (75×25 mm). (b) Chemotaxis experiment, t=6 h. Chamber B is filled with a MNP suspension (rusty discolouration). *P. caudatum* cells were microscopically observed to have migrated from chamber A→B.

Initial control experiments designed to determine the rate of transfer of both inanimate microparticulates and *P. caudatum* cells between each chamber were designed as follows. For *P. caudatum* experiments, approximately 25 cells were transferred to chamber A in 100 *μ*l of DTW. The EEs were placed on the stage of a stereomicroscope which was focused on chamber B. The sample was kept static for the duration of the experiment and was observed regularly every 4-6 hours. The microscope’s halogen lamp was switched off in between observations and the entire setup was exposed to a day/night cycle whilst being kept out of direct sunlight.

Experiments examining particulate diffusion were conducted as above but the EEs had a concentrated amount of exogenous particles added to chamber A rather than cells. Two varieties of particulate were added:

- 2.0 *μ*m diameter carboxylate-modified latex microspheres labelled with fluorescein (Sigma Aldrich, Germany) (hereafter, FLPs, ‘fluorescent latex particles’). 100 *μ*l; of stock solution (2.5% solids, approx. 7.2×10^9^ particles per ml) was added to give a total concentration for chamber A of 0.125% solids w/v (approx. 7.20×10^8^ particles).
- 200 nm diameter multi-core magnetite (iron II/III oxide) nanoparticles, prepared with a hydrodynamic starch coating (Chemicell, Germany) (hereafter, MNPs, ‘magnetite nanoparticles’). 100 *μ*l of stock solution (25 mg/ml) was added to give a total concentration for chamber A of 1.25 mg/ml (approximately 2.75×10^11^ particles).

These two varieties of particulate were chosen for these and consequent experiments as our previous work has demonstrated that *P. caudatum* will favourably ingest them whilst suffering no apparent deleterious health effects [31, 26]. Samples were taken every 4-6 hours by simultaneously drawing 10 *μ*l of fluid from both chambers and examining the sample from chamber B with a Zeiss Axiovert 200M inverted microscope; samples were checked at 200 × magnification with fluorescence microscopy in the case of the FLPs and 600-1000 × using phase contrast optics for the MNPs. Although the particle size of the MNPs was beyond the resolution of the light microscope, they are known to flocculate into micro-scale quantities in the presence of salts. A magnet was also held close to the edge of the droplet in order to observe contrails and discolouration in the fluid due to the movement and concentration of magnetite in areas closer to the magnet.

### 1.3. Controlled transport of particulates

Two methods of controlling *P. caudatum* migration between the two chambers were investigated, one via attraction and the other repulsion.

#### 1.3.1. Chemoattraction

Firstly, attraction of *P. caudatum* between chambers was investigated by simultaneously transferring approximately 25 cells into chamber A in 100 *μ*l of DTW and the same volume of one of three varieties of *P. caudatum* ‘food’ into chamber B. Thus the hypothesis that *P. caudatum* cells would migrate from chamber A to B, guided by a chemoattractant gradient (chemotaxis). The three varieties of food were as follows:

1. *Saccharomyces cerevisiae* (yeast). Yeast were cultured from approximately 20 grains (0.1 g) of freeze dried, commercially-available bread yeast (Allinson, UK) in 10 ml of DTW containing 0.1 g of glucose in an incubator at 22*°*C, for 1 hour. The cultures were then transferred into boiling tubes and placed in a 100*°*C water bath for 15 minutes. A few drops of 40% Congo red dye dissolved in ethanol were added before cultures were reserved in a refrigerator until use.
2. FLPs, 100 *μ*l of stock solution to bring total concentration to 0.125% (w/v), approx. 7.20 × 10^8^ particles.
3. MNPs, 100 *μ*l of stock solution to bring total concentration to 1.25 mg/ml, approx. 2.75 ×10^11^ particles.

Chemotaxis experiments were performed in a similar manner to those described in section 1.2: the EE was placed onto a stereomicroscope stage and the was observed at a position over chamber B once per hour for the presence of *P. caudatum* cells. When cells were found to have migrated to chamber B, samples were taken by drawing off the cuvette’s fluid and adding it to an equal quantity of 4% paraformaldehyde in pH 7.2 phosphate buffered saline (PBS). Individual *P. caudatum* cells were isolated under a stereomicroscope, transferred to a cavity slide on the inverted microscope and checked optically for red-stained yeast or MNPs, or for FLPs with fluorescence.

A further experiment was designed in order to assess whether multiple varieties of particulate could be collected by *P. caudatum* cells, towards describing their ability to perform parallel manipulation. Another set of chemotaxis experiments was conducted where 100 *μ*l of a 50:50 mixture of stock (see above) FLPs and MNPs were added to chamber B; following the successful migration of cells to chamber B, they were checked for the presence of both varieties of particle.

#### 1.3.2. Galvanorepulsion

Repulsion experiments were performed through the use of a DC electrical field, along the hypothesis that *P. caudatum* migrates towards the cathode in a system where a small electrical current is injected (galvanotaxis) [32]. Two 90×1.0 mm platinum electrodes were inserted into the EE and connected to a benchtop DC power supply providing 18 V at a maximum current of 0.3 A. Circa 25 *P. caudatum* cells were introduced into chamber A of the EE, along with the anode, in 100 *μ*l of DTW. The cells were allowed 5 minutes to acclimatise to their new environment before the power supply was switched on. The experiments were observed continuously using a stereomicroscope and recorded using a Brunel Eyecam (Brunel Microscopy, UK). Experiments were repeated 5 times for each chamber being observed (i.e. 5 at A, 5 at B).

### 1.4. Controlled retention and deposition

#### 1.4.1. Retention

Retainment of particulate cargoes over the duration of EE experiments was investigated by transferring 5 *μ*l of concentrated *P. caudatum* culture to a clean culture vessel containing 0.015% w/v solution of FLPs (approx. 5.4×10^7^ particles). Cells were left for 1 hour before being manually removed with a micropipette and placed in 5 *μ*l of fresh media. This was achieved by transferring 200 *μ*l of the original culture media to a large cavity microscope slide containing a drop of quieting solution (1% methyl cellulose), which slowed the organisms’ migration sufficiently to manually collect them in 2.5 *μ*l of media using a micropipette. This step was essential in order to ensure that no extracellular particulates were transferred into the fresh medium. Five cultures were run in parallel such that one could be examined each hour for 4 hours and the final one was examined after 24 hours: examination included pipetting the entire volume onto a series of microscope slides and observing for exogenous FLPs under fluorescence optics. Each set of retention experiments were run in triplicate.

#### 1.4.2. Deposition

The following ‘deposition’ experiments were designed to demonstrate the principle that *P. caudatum* cells loaded with exogenous particles could be made to cease all movement (transport operations) after a specific point using a simple and effective method, which led to the deposition of their cargoes in a predictable manner. This was achieved by transferring *P. caudatum* cells containing FLPs, which were prepared in the same manner as in the aforementioned ‘retention’ experiments, to chamber A in an EE. 200 *μ*l of 4% paraformaldehyde in PBS was added to chamber A in order to fix all of the resident cells, whilst 200 *μ*l of DTW was added to chamber B. Both a stereomicroscope and fluorescence inverted microscope were used to observe the distribution of fixed cells in chamber A, after which the fluid was carefully drawn off and observed as per the ‘retention’ experiments for evidence of extracellular FLPs. Experiments were repeated in triplicate.

### 1.5. Programmability

‘Programming’ of EEs with multiple input types was investigated via a chemoattraction operation followed by galvanorepulsion. Approximately 650 *P. caudatum* cells in 200 *μ*l of DTW were placed into chamber A and a corresponding volume of a particulate solution (100 *μ*l of both MNPs and FLPs at stock concentration) was delivered into chamber B simultaneously: the experiment was run as per the chemotaxis experiments in section 1.3.1, with the exception that a platinum cathode was placed in chamber A and an anode in chamber B, as per section 1.3.2. After 6 hours had elapsed and cells had been positively identified as being present in chamber B with a stereomicroscope, the electrodes’ power supply was switched on. Videomicrography was started at the point in the experiments where the electrodes were turned on. Experiments were repeated 10 times.

### 1.6. Automation

The principle of detection of FLPs via fluorescence spectroscopy was chosen to be the simplest and lowest cost method for detecting the completion of a transport operation. A fluorescence spectrometer designed to articulate onto a single EE chamber was designed as follows (see Supplementary Information 3 for parts list).

The light source used was a single surface mount 485nm light emitting diode (LED) that produced 479 Lux. The LED was mounted to a custom board with a large aluminium heat-spreader, onto which a 15*°* collimator lens was affixed in order to focus the light produced. The LED was driven by a benchtop power supply at 3.1 V and 250 mA, switched via a relay controlled by an Arduino Uno microprocessor board based on the ATmega328 platform (Arduino, Italy) (microprocessor code is included in Supplementary Information 2). The LED board and lens assembly was mounted to an EE cuvette via a slip-on 3D printed adapter (Fig. 2a), which was generated in OpenSCAD on an open source template [33]. The adapter, which fit snugly around three sides of the cuvette (Fig. 2b), also served to mount two filters and a light dependent resistor (LDR) at a 90*°* angle to the light path (Fig. 2d-e). The filters used were an excitation bandpass filter (i.e. between the light source and sample) and an emission notch filter (between sample and detector); inspired by Ref. [34], we used low-cost photography lighting gels as filters.

**Figure 2:**
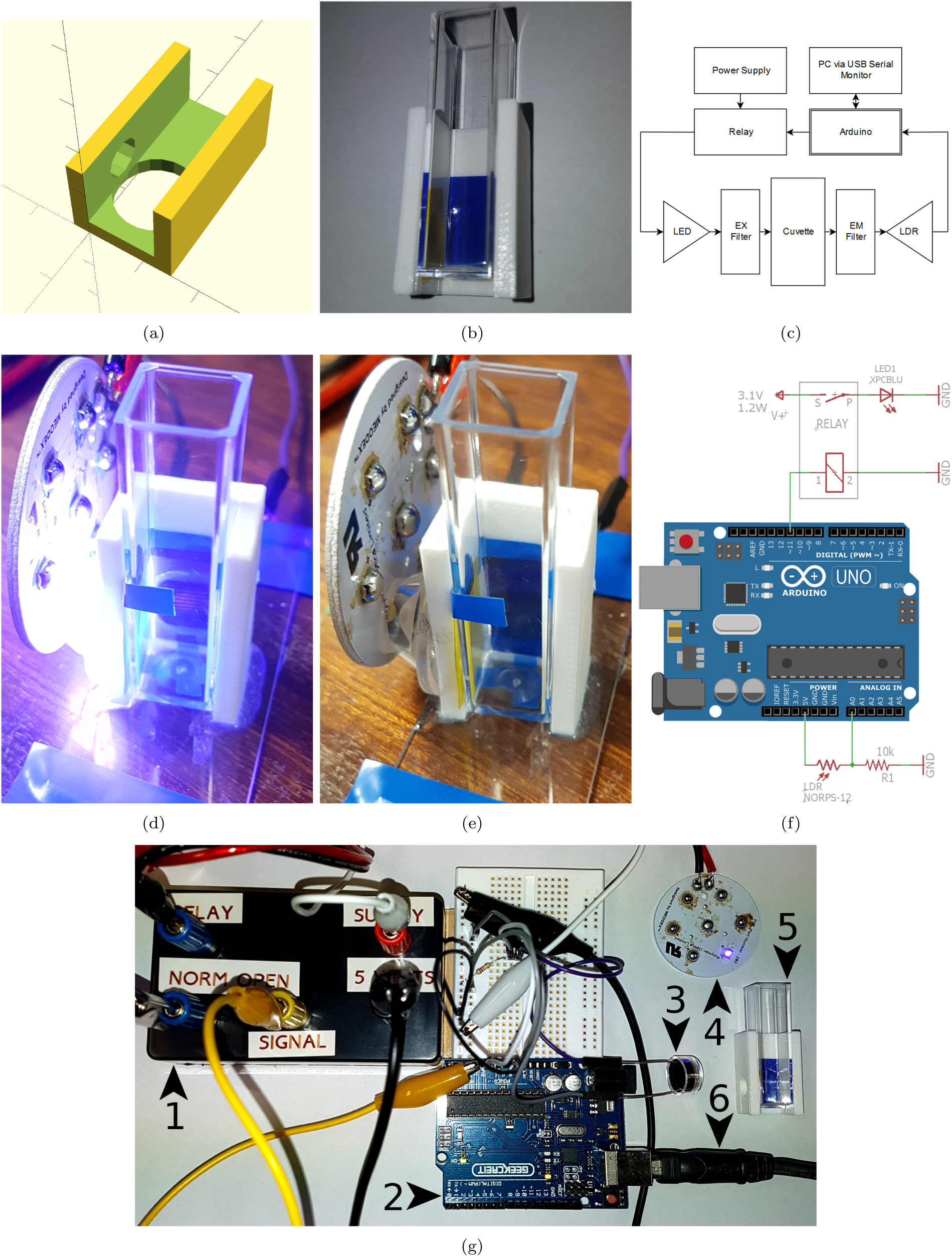
EE fluorescence spectrometer. (a) 3D printed cuvette adaptor. (b) Photograph of cuvette adapter with filters and cuvette *in situ*. (c) System diagram. (d-e) Photographs of assembled LED with heatsink, LDR (rear), collimator, adapter and cuvette, with and without LED illumination. (f) Wiring diagram. (g) Partially assembled spectrometer system, where (1) relay, (2) Arduino, (3) LDR, (4) LED, (5) cuvette, filters and adapter, (6) USB connection.

The light intensity was measured by the LDR, using a voltage divider with a resistor (10 kΩ in this instance), with the output of the voltage divider read by an analogue pin on the Arduino microcontroller. The on-board 10-bit ADC converted the analogue signal into a digital value which was sent to the PC via serial monitor USB link. The system is illustrated in Fig. 2; (c) system schematic, (f) wiring diagram, (g) partially-assembled.

The operation cycle of the spectrometer was as follows: the LED is switched on for 1 second and the LDR value is read and printed on-screen. The LED then switches off for 1 second and the LDR value is printed again. The first number returned gives the relative sensor value, indicating the quantity of fluorescent compound in the sample. The second number indicates a reference value as an internal control for each reading.

The spectrometer was always operated inside an opaque box in order to eliminate interference by ambient light. Initial device evaluation was performed on graduated series of FLPs, inside a single cuvette.

Further experiments were performed with the spectrometer *in situ* on chamber A of EEs following experiments identical to those detailed in section 1.5, with the exceptions that only FLPs were added (200 *μ*l of stock solution), cells were only exposed to the particles for an hour and a range of cell densities was investigated. The rationale behind this experiment was that the spectrometer would detect when cells had returned to chamber A following being ‘programmed’ to return there after internalising a quantity of FLPs. The spectrometer was used 5 minutes following the application of the DC field. A small number of cells from each experiment were withdrawn, fixed and imaged in order to estimate the average number of FLPs within each cell.

## 2. Results

### 2.1. Suitability of experimental environment

In initial control experiments, it was found that the organisms did not migrate to chamber B within, at minimum, the first 24 hours of observation. Even so, only a few of the 25 initial *P. caudatum* cells were observed in chamber B after 48 hours in all experiments (maximum 4).

No FLPs or MNPs were observed to have diffused to chamber B over the 48 hour experiments, presumably due to both varieties of particle not diffusing as a product of their size and/or density. Data from these experiments are shown in Supplementary Information, section Appendix A.3.

### 2.2. Controlled transport

#### 2.2.1. Chemoattraction

*P. caudatum* cells were observed to migrate from chamber A→B in shorter time scales than control experiments when guided by all three attractants. The average time for the organisms to migrate to the supplied attractants were: yeast 1.6 h, FLPs 5.4 h, MNPs 1.8 h (Fig. 1b), FLPs and MNPs 1.8 h (see Appendix A.4 for dataset). On microscopic examination of fixed cells recovered from chamber B after 24 hours of exposure to each attractant source, all three varieties of particle could be distinguished in the cells’ cytoplasm. Results therefore suggest that all varieties of particulate are effective chemoattractants. Furthermore, cells that were exposed to FLPs and MNPs simultaneously were observed to have ingested quantities of both (Fig. 3), indicating the *P. caudatum* may be loaded with multiple particle types for parallel operations

**Figure 3:**
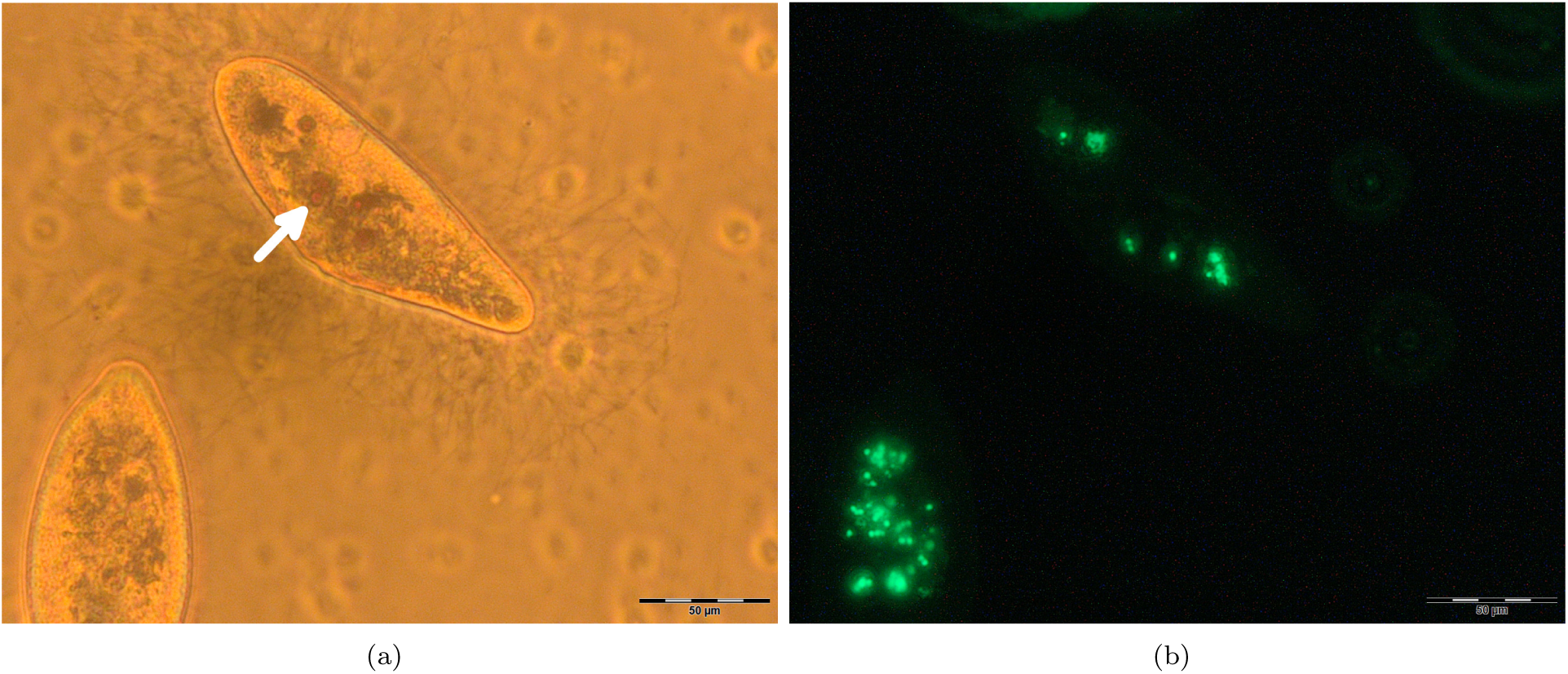
Photomicrographs to show fixed *P. caudatum* cells following exposure to a 50:50 mixture of FLPs and MNPs. (a) Brightfield. Rust-coloured intracellular inclusions indicative of magnetite clusters are present (arrowed). (b) Fluorescence. Fluorescent intracellular objects consistent with the FLPs are also present.

#### 2.2.2. Galvanorepulsion

*P. caudatum* cells were observed to respond to the application of a DC field by immediately altering their swimming direction towards the cathode in chamber B (Fig. 4, Supplementary Information File: SI Movie 1). Despite their rapid response, the cells required up to 5 minutes of constant stimulation to evacuate their chamber due to their slow speed relative to the dimensions of the EE and their directional migration being vague, i.e. cells would frequently collide with the walls of the cuvette and the sides of the connecting tube several times before finding the correct path. There was frequently a small number (range 0-4) of cells that did not move towards the cathode within the 5 minutes time frame and occasionally a few cells would stay in the connecting tube rather than emerge into chamber B.

**Figure 4:**
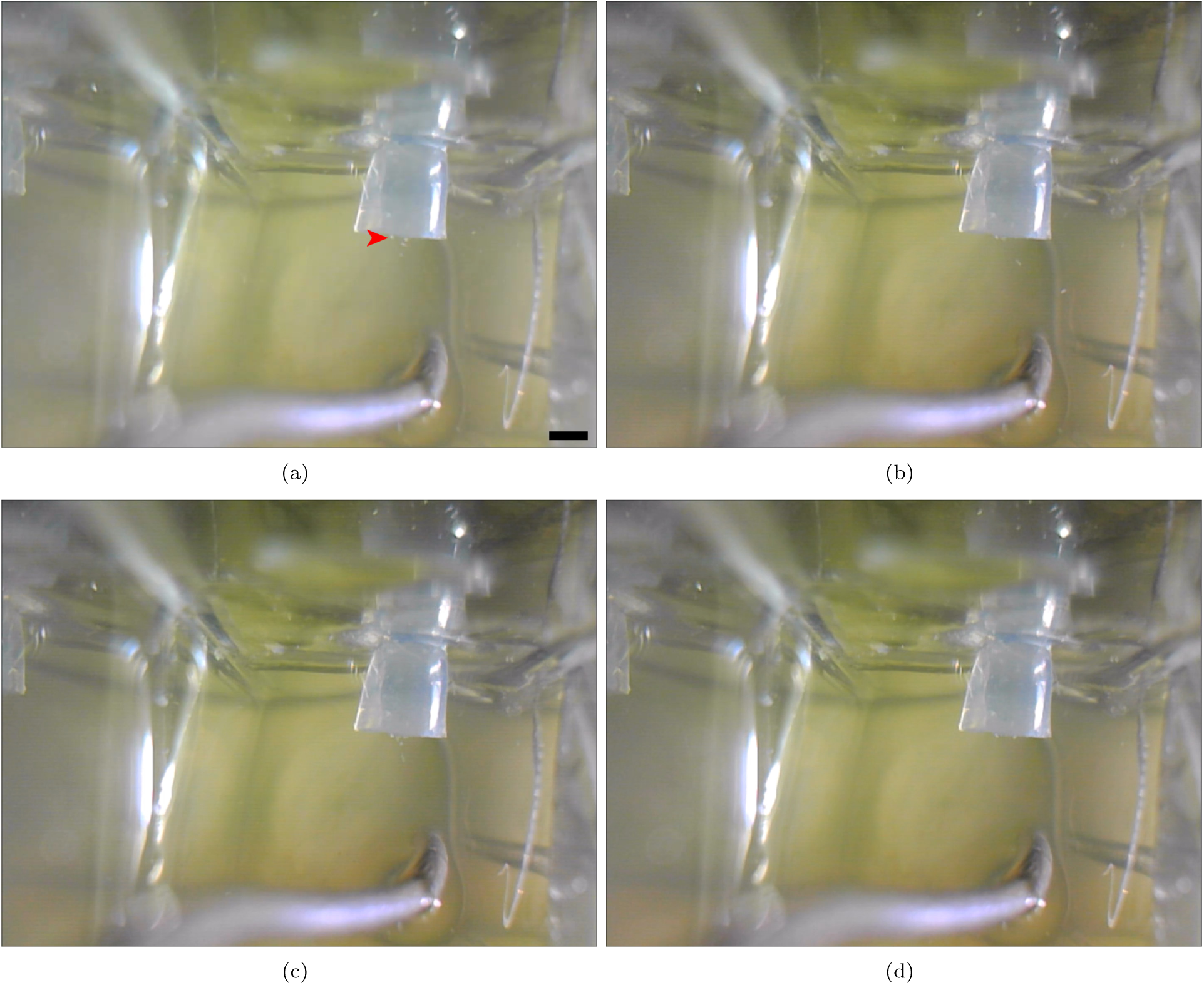
Stereomicrographs to demonstrate the movement of *P. caudatum* cells away from a live anode (silver object in bottom central third of images). Several cells (cluster arrowed in [a]) can be seen swimming towards, then down, the linking tube that connects the chambers in the EE. Images interval 5 seconds. Scale bar in [a] 1 mm.

### 2.3. Controlled retention and release

Extracellular FLPs were not identified in culture media until at least 24 hours had elapsed. Furthermore, organisms were observed to retain FLPs for up to 4 days post-exposure (data not shown), indicating that the organisms do not excrete them for the duration of their lifespan.

Fig. 5 shows the results of a representative experiment examining the deposition of fixed cells in an EE chamber: cells were found to fix rapidly and remain dispersed throughout the chamber, rather than settle to the bottom. Fluorescence microscopic examination of the culture media revealed no evidence of particles being released from cells as a result of fixation (i.e. autolysis was unlikely to have occurred).

**Figure 5:**
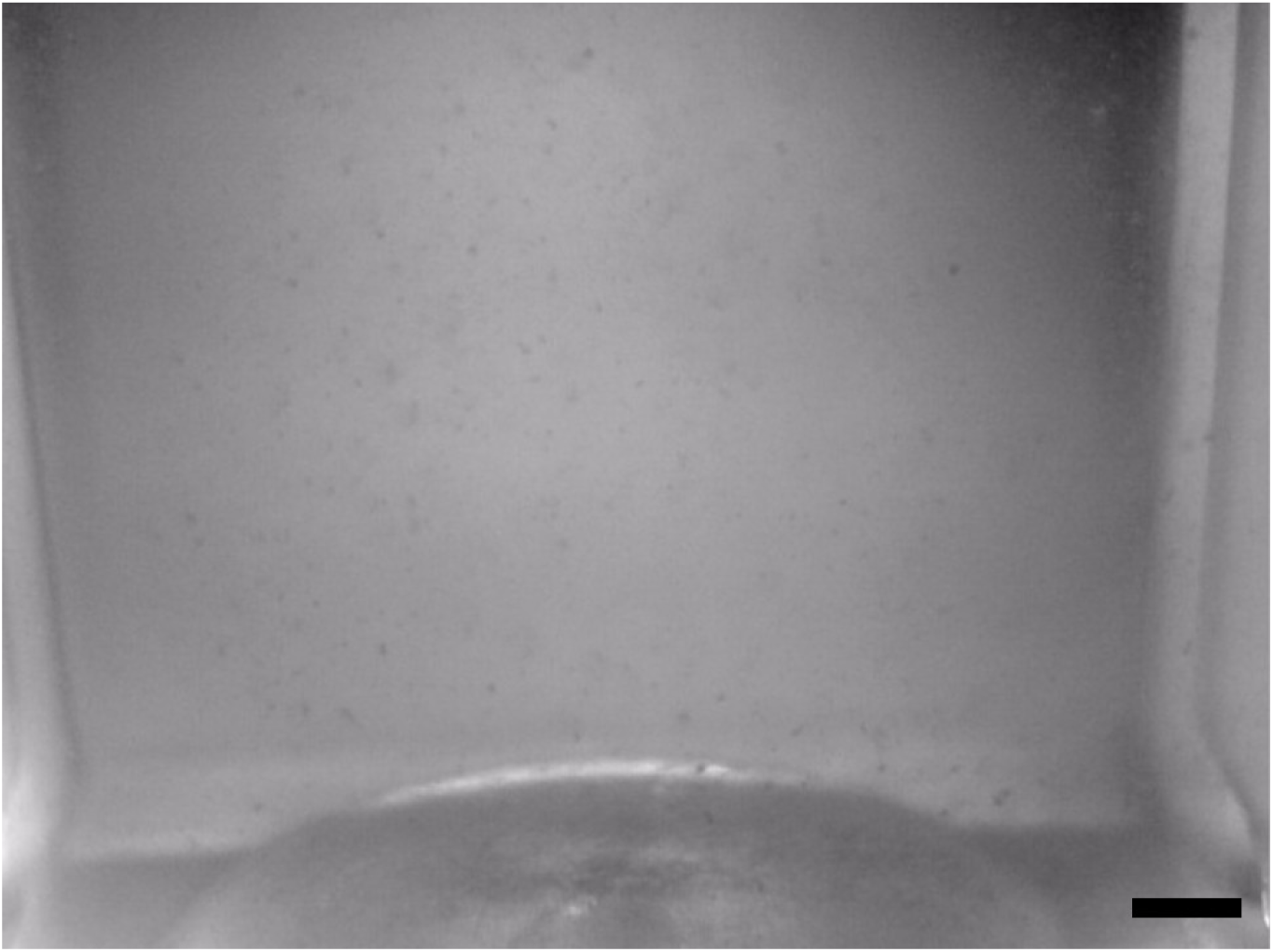
Photograph to show distribution of fixed *P. caudatum* cells inside an EE chamber. The cells have the appearance of particulates. Scale bar 1 mm.

### 2.4. Programmability

*P. caudatum* cells responded to multiple concurrent inputs as intended: cells were observed to migrate from chamber A→B in the manner described in chemotaxis experiments and consequently B→A when the DC field was applied. The number of cells that remained in the anodic chamber were similar to those seen in the galvanorepulsion experiments; again some cells (range 2-12) remained in the anodic chamber and/or remained within the link tube, but the majority of organisms migrated back towards the cathodic chamber in all experiments.

### 2.5. Automation

The fluorescence spectrometer device was found to be sensitive to a minimum of quantity of approximately 4.5 ×10^4^ FLPs; increases or decreases in this quantity of particle equated to, at the least sensitive range of the sensor’s operation, relative changes of 8.33 in a total range of 1024 (s.d. 1.24) (see Supplementary Information 3 for calibration dataset). A calibration curve for the spectrometer is shown in Fig. 6a and shows that the sensor’s response was not linear, possibly due to the sensor beginning to saturate at higher concentrations of FLPs.

**Figure 6:**
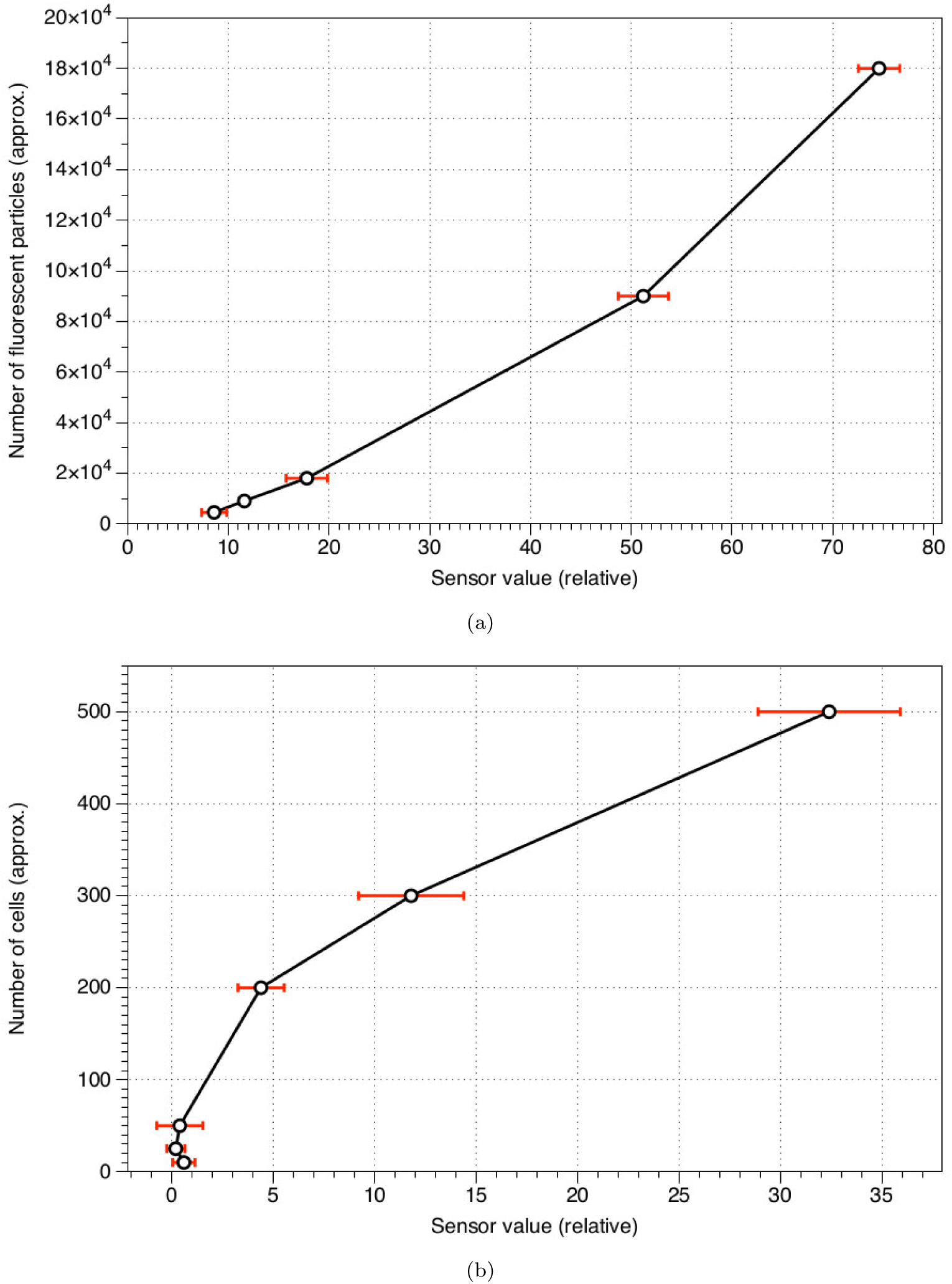
Charts to show spectrometer output. Error bars to 1 S.D. (a) Graduated concentration of FLP solution. (b) *P. caudatum* cells (per ml) loaded with FLPs.

The spectrometer was less sensitive with regards to detecting cells loaded with FLPs but was able to detect a minimum of 200 cells per ml (i.e. approximately 400 organisms total) (Fig. 2b). Cells that were examined post experiment revealed that the number of particles contained within each cell was highly variable, but was on average 161 (standard deviation 86, range 48-345) per cell (Fig. 7) (see Supplementary Information 3 for dataset).

**Figure 7:**
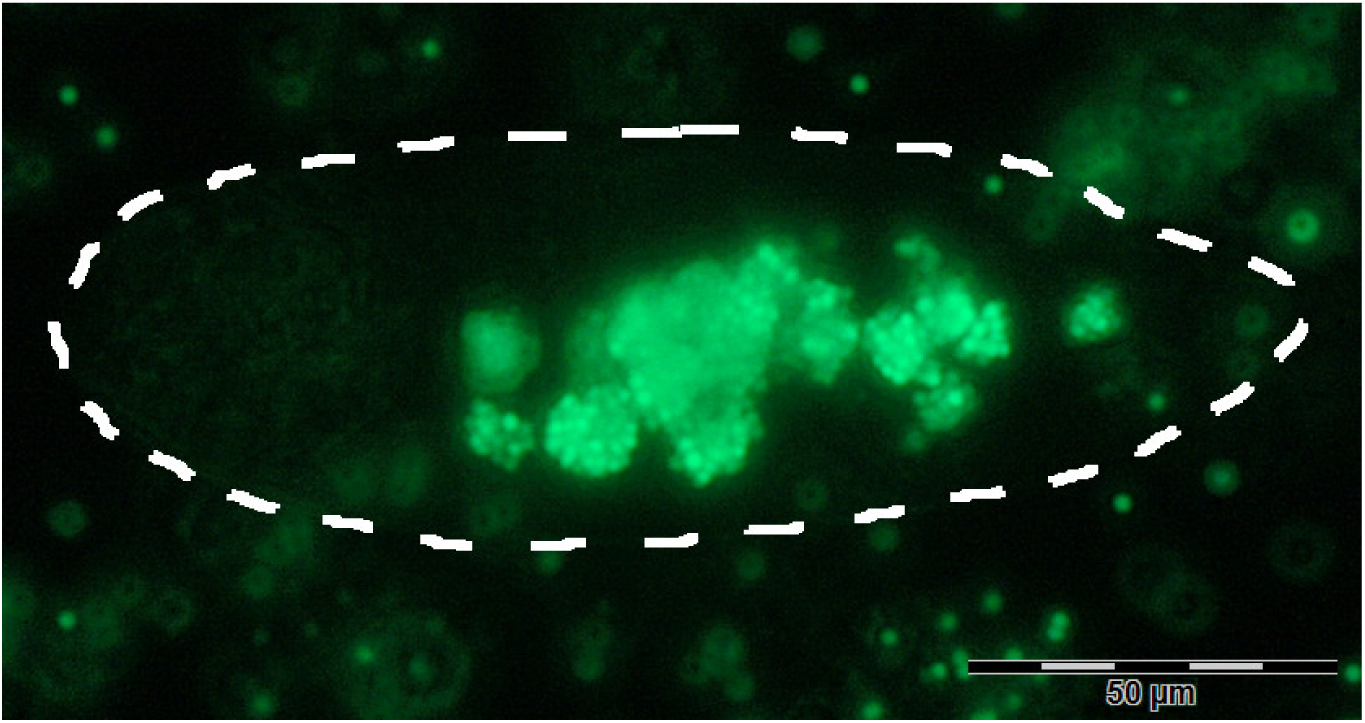
Fluorescence micrograph to show clusters of fluorescent particles inside intracellular vesicles of a fixed *P. caudatum* cell. The outline of the cell is highlighted.

## 3. Discussion

### 3.1. Evaluation of devices

The basic EE was reasoned to be suitable for use as *P. caudatum* cells did not migrate freely between chambers within the time frame of device operation (<36 hours) without additional stimuli. Factors such as nutrient abundance in their original chamber and the confined geometry of the environment causing spontaneous alternation behaviour likely contributed to the organisms’ propensity for not rapidly exploring their entire environment.

Observations on the amount of time *P. caudatum* took to migrate between chambers indicated that the presence of ‘food’ (yeast, MNP starch coatings) may dramatically increase the rate of the organisms’ migration between chambers; FLPs did not attract as strongly, despite their being eaten *en masse.* Particulate coating is therefore an attractive route towards increasing the rate of transport operations.

Galvanorepulsion was found to be a quick and effective method for controlling *P. caudatum* migration, but it suffered drawbacks stemming from the issues associated with passing a current through highly resistive fluid media. Significant ‘fine tuning’ was required in order to increase the power output to a level that would be effective across the length of the device without causing cells to rupture spontaneously in proximity to the electrodes. This phenomenon need not be a detriment in future iterations of such devices, however, as it could be used as a method of inducing particulate decellularisation and deposition, i.e. the cells are killed once they have completed migration.

*P. caudatum* retains ingested particulates long enough (> 24 hours) for transport operations to complete. Some progress was made towards controlled deposition, although the method used (chemical fixation) leaves cells suspended randomly in solution, does not release particulates from their cells and contaminates the EE. Furthermore, fixation precludes the use of fluroescence spectrometry after the deposition stage due aldehyde exposure inducing autofluorescence in biological tissues.

Regarding device programmability, whilst we demonstrated that the use of multimodal input is sufficient to complete several operations in one device, there is little room for continuous reprogramming or cascading of operations due to several processes (chemoattraction, deposition via killing the cells) being undynamic. Future work focusing on more dynamic stimuli, such as light, may therefore be productive.

The spectrometer was found to detect FLPs in minimum concentrations of approximately 50,000 and exceeding 180,000 in 2 ml of fluid. In comparison with the sensor readings for *P. caudatum* cells loaded with the FLPs, the minimum number of cells detected was 200 per ml (400 total): the mean relative sensor values for 200, 300 and 500 cells per ml were 4.4, 11.8 and 32.4 respectively, which equates to approximately 100, 160 and 500 particles per cell, according to the calibration curve for free particles. Microscopic observations of cells treated with FLPs, revealed that they ingest and retain highly variable quantities, indicating that calibration of the spectrometer against microscopically-determined values is essential for quantitative measurements. Factors such as uneven distribution of cells in their chambers, cell division:death ratios, cell density-mediated feeding rates and various fluorescence artefacts (autofluorescence, quenching etc.) all likely contribute to the discrepancy between calibration and experimental results.

Based on microscopic observations, we estimated the average mass transfer of FLPs by a *P. caudatum* cell to be about 14 mg/hour (0.68 g in 6 hours by 1000 cells in 4 ml total fluid volume) over a 5 mm distance, although this value is subject to significant error due to the variation in particle quantities ingested between organisms. Our other results indicate that this rate may be significantly increased if the particulates are made more ‘palatable’ for the cells.

### 3.2. Device development and applications

Excluding hardware, optional components and the platinum wires (which could likely be replaced by cheaper alternatives), the unit cost for a single EE/spectrometer was approximately 65 GBP. This price was significantly cheaper than all commercially-available spectrometers we found via an internet search. Adaptations such as introduction of a second LED/filter set and optimising LDRs to specific wavelengths would be cheap and could increase the sensitivity and range of uses for these devices.

Further adaptations for increasing the ‘usefulness’ (modularity, scalability, parallelism etc.) of the devices presented here could include: further detectors, e.g. metal detection via a Hall effect sensor for metallic particulates or electrical capacitance measurement for cell movement; automated killing/lysis of cells via a microcontroller-driven chemical pump or large electrical current; use of further dynamic input types such as light and temperature.

We conceive the principle applications of cilia-mediated particle manipulation devices such as these to be in adaptive transport of microparticulates, especially in industries such as environmental clearance of polluting nano- and microscale inorganic objects (taking into consideration *P. caudatum’s* demonstrated tolerance to certain nanomaterials [31]), as well as micromixing/microfluidics. There is also scope to interpret these devices in terms of unconventional computing: if a third chamber (C) were introduced and input were applied to both A and B, the ‘output’ could be interpreted in terms of a logical TRUE if particulates are (or are not) delivered to C. Such a configuration could be interpreted as several varieties of gate (e.g. OR, NANO) driven by an automated bio-computer interface.

## Acknowledgements

The work was supported by the Leverhulme Trust grant “Towards Artificial Paramecia” (grant number RPG-2013-345).

## Appendix A. Supplementary Information

### Appendix A.1. Key to SI files

**SI 1**: Video to show *P. caudatum* cells in an EE chamber, next to a platinum anode. The cells may be observed to rapidly enter the linking tube and migrate in the direction of the other chamber, which contains the cathode.

**SI 2**: Arduino sketch used for DIY fluorescence spectrometer.

**SI 3**: Spreadsheets showing spectrometer parts list, control experiments dataset, chemotaxis experiments dataset, spectrometer calibration data and intracellular fluorescent particle counts.

### Appendix A.2. Estimation of diffusion rate between EE chambers

Simple diffusion between both chambers was estimated to take a minimum time of approximately 3 hours 30 minutes: by 3Equation A.1

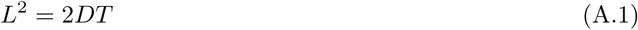

Where L is length, D is the coefficient of diffusion (estimated at maximal rate of 10^−9^ in liquids) and T is time, thus the time to diffuse 5 mm would be 12,500 seconds (208 minutes).

### Appendix A.3. Results of Control Experiments

Results from initial experiments described in section 2.1 measuring the rate of diffusion of (a) *P. caudatum* cells between the two chambers in an absence of stimuli and (b) particulates between the two chambers are shown in Supplementary Information 3.

### Appendix A.4. Results of Chemotaxis Experiments

Dataset in Supplementary Information 3 shows results referenced in section 2.2.1, indicating the time taken for *P. caudatum* cells to traverse the linking tube between chambers in their EE to begin feeding on the supplied particulate ‘foods’.

### Appendix A.5. Estimation of transferred particulate mass

The approximate mass of a single latex particle was calculated by the mean volume of a 2 *μ*m latex sphere multiplied by the manufacturer-specified density, is shown in equation A.2.

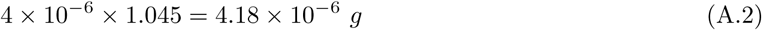

Multiplied by the average number of particles per cell as reported in section 2.5, 163 (range 48-345), we estimate the mass transfer of FLPs by *P. caudatum* cells to be 6.81 × 10^−4^ g (2.00 × 10^−4^-1.44 × 10^−3^g) per operation, over 6 hours (as per the experiment detailed in section 1.5). This equates to an estimated total mass transfer of 0.68 g (0.20-1.44 g) for the maximum cell densities that were tested (1000 in 2 ml).

